# Extracellular Ca^2+^-Sensing Receptor (CaSR) Regulates Hypothalamic Function to Control Energy and Skeletal Metabolism in Mice

**DOI:** 10.1101/2025.05.22.655605

**Authors:** Jennifer Park-Sigal, Mariana Norton, Chia-Ling Tu, Zhiqing Cheng, Nicole Fadahunsi, Hokyou Lee, Sunhee Kim, Alfred Li, Lea T Grinberg, Gavin A Bewick, Kevin G Murphy, Wenhan Chang

## Abstract

The extracellular Calcium-sensing receptor (CaSR) regulates cellular responsiveness to physiological changes in ionized calcium (Ca^2+^) concentrations. The CaSR is expressed in the brain, including in hypothalamic growth hormone stimulating GHRH and anorectic POMC neurons that control growth and energy homeostasis. We embryonically deleted the *Casr* gene in neurons to create ^Neuron^CaSR^-/-^ mice to delineate the role of this receptor in regulating growth, skeletal development, and energy metabolism. ^Neuron^CaSR^-/-^ mice had reduced size, weight and bone mass compared to littermate controls, with a dysregulated growth hormone axis. They also showed increased adiposity and circulating leptin levels, leptin resistance, and decreased glucose tolerance, along with reduced expression of the anorectic precursor peptide POMC and secondary increases in the expression of the anorectic peptide AgRP in the hypothalamus of ^Neuron^CaSR^-/-^ mice. Knockdown of CaSR in adult mice specifically in the hypothalamic arcuate nucleus, where GHRH, POMC and AgRP neurons reside, also resulted in increased body weight, adiposity, leptin resistance, and glucose intolerance, and reduced bone mass. Together these data suggest that neuronal CaSR critically regulates energy and skeletal metabolism and body growth by modulating hypothalamic function, representing a new paradigm for central integration of calcaemic activities with body function.

## Main Text

Calcium (Ca^2+^) is essential for all cell functions. The extracellular calcium-sensing receptor (CaSR), a member of the family C G-protein couple receptors (GPCRs), has a well-defined role in systemic Ca^2+^ homeostasis, but is widely expressed in tissues without apparent roles in systemic Ca^2+^ regulation, including the brain^1,2^. However, the role of neuronal CaSR in general physiology remains largely unknown.

CaSR was first identified in parathyroid cells (PTCs) where it responds to changes in extracellular [Ca^2+^] ([Ca^2+^]_e_) by adjusting PTH secretion to maintain systemic mineral and skeletal homeostasis^3^. Inactivating mutations of the *CASR* gene lead to neonatal severe hyperparathyroidism (NSHPT)^2,4–6^. These patients manifest high serum PTH and Ca^2+^ levels, severe growth retardation, and de-mineralized skeletons^5,7^. Similar developmental defects have been shown in ^Global^CaSR^-/-^ mice in which the exon 5 of the *Casr* gene was deleted in germline cells^8^. Although PTH excess has been assumed to drive the mineral defects, whether and how reduced CaSR activities in extra-parathyroid tissues cause growth and skeletal defects remains to be defined.

The CaSR is also expressed widely in the brain^1^. In cultured neurons, raising [Ca^2+^]_e_ activates neuronal ion channels, promotes neurite outgrowth, enhances neuronal migration and proliferation, and can increase gonadotropin-releasing hormone (GnRH) secretion, potentially by activating the CaSR^1^. These *in vitro* data indicate the ability of neurons to respond to changes in the local and perhaps the systemic Ca^2+^ environment. Neuronal CaSRs may therefore be involved in the central control of skeletal and energy metabolism in response to changes in Ca^2+^ availability in the body, as Ca^2+^ deficiency can retard growth and skeletal development and cause metabolic imbalance^9^. However, such roles of neuronal CaSR in the central nervous system (CNS) *in vivo* has not been determined.

The hypothalamus integrates autonomic nervous signals and releases hypothalamic releasing factors to modulate the secretion of hormones from the anterior pituitary essential for growth, reproduction, energy homeostasis, and skeletal development^10–13^. Specifically, the arcuate nucleus (Arc), located at the base of the hypothalamus, contains growth hormone releasing hormone (GHRH) neurons, and also anorectic neurons that express pro-opiomelanocortin (POMC), which reduce food intake and increase energy expenditure, and orexigenic neurons that express both agouti-related protein (AgRP) and neuropeptide Y (NPY), which increase food intake and suppress energy expenditure^14–16^. GHRH, POMC and AgRP/NPY neurons respond to leptin, a protein hormone produced by white adipocytes as a signal of energy storage, and to other hormonal factors and circulating glucose to balance energy metabolism and growth^16–18^. Together, POMC and AgRP/NPY neurons constitute a feedback mechanism to homeostatically maintain body weight and body fat. In POMC neurons, activation of the leptin receptor (LEPR) stimulates Signal Transducer and Activator of Transcription 3 (STAT3)-mediated signaling cascades to promote the secretion of α-melanocyte-stimulating hormone (α-MSH)^19^ that produces anorexigenic actions *via* the activation of the melanocortin 4 receptor (MC4R)^16,20^. In contrast, LEPR activation suppresses the production and release of AgRP and neuropeptide Y (NPY) in AgRP neurons^21,22^, blocking the anorexigenic effects of α-MSH and driving hunger through Y1 and Y5 receptors. Obesity results in a relative insensitivity to leptin signaling, and leptin insensitivity in POMC and AgRP neurons leads to obesity and diabetes in patients and animal models^15,23^. However, leptin has also been identified as a key hormone in maintaining bone mass^24^, and dysregulated leptin responses in the CNS have been shown to contribute to the development of skeletal disorders in patients and animal models^25^.

There is thus increasing interest in the mechanisms that integrate the central regulation of skeletal and energy metabolism in response to changes in Ca^2+^ homeostasis. We investigated the role of neuronal CaSR in the coordination of skeletal development and body growth, by studying a neuron-targeted CaSR knockout (^Neuron^CaSR^-/-^) and Arc-specific CaSR-knockdown mouse models.

### CaSR expression in the hypothalamus and generation of neuron-specific CaSR knockout (^Neuron^CaSR^-/-^) mice

We detected CaSR protein in >70% of neurons by immunohistochemistry in the hypothalamic Arc, where it was localized to the cell membrane and intracellular stores of GHRH- and POMC-expressing neurons, and to a much less extent in AgRP-expressing neurons, which were concurrently marked with corresponding fluorescent RNAScope probes, in mice and humans (**Fig. 1A,B, Extended Data Fig. 1**). To determine its functions *in vivo*, we made ^Neuron^CaSR^-/-^ mice by breeding CaSR^fl/fl^ mice^3^ with mice expressing Cre recombinase under the control of a rat *Nes* (Nestin) gene promoter^26^. ^Neuron^CaSR^-/-^ mice were born in a normal Mendelian ratio and survived to all time points of analysis. Genomic DNA analyses demonstrated that the Cre-mediated recombination of *Casr* gene (i.e., deletion of exon 7) occurred in the brain, but not in the pituitary, bone, or parathyroid gland (**Fig. 1C**). Slight gene recombination, indicated by a faint 285-bp DNA band, in lung, intestine, and kidney, was surmised to reflect the existence of a small number of neuronal elements and Nestin-expressing stem cells in those tissues.

**Fig. 1.**
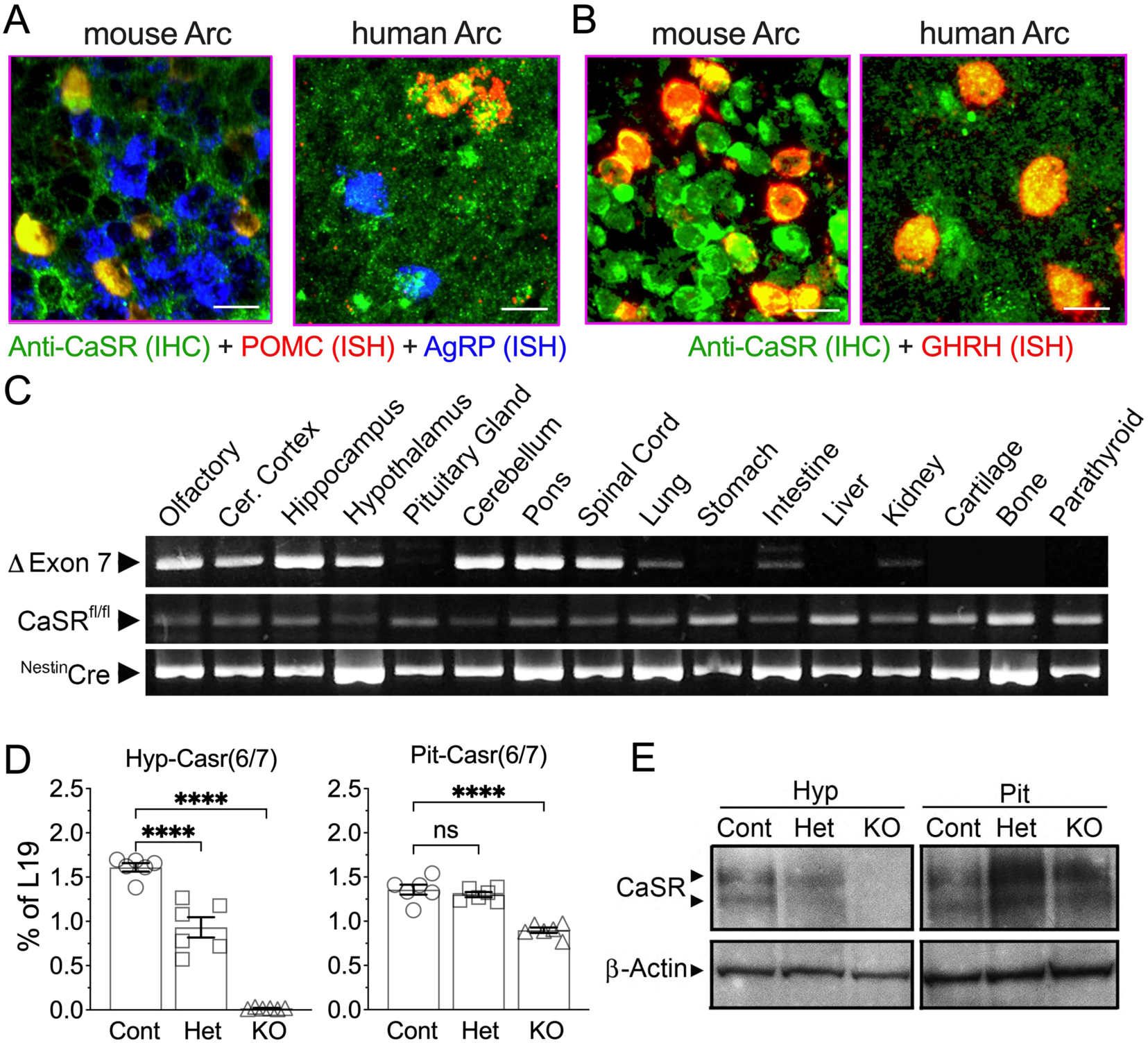
Expression and localization of CaSR in POMC and GHRH neurons in mouse and human hypothalami and ablation of CaSR gene in the CNS of ^Neuron^CaSR^-/-^ mice. (**A,B**) Immunohistochemical (IHC) detection of CaSR (in green) with concurrent in situ hybridization (ISH) of POMC (in red) and AgRP (in blue) RNA in the panel **A** or GHRH RNA in the panel **B** show localization of CaSR in POMC- and GHRH- expressing neurons (in yellow), but to a much less extent in AgRP-expressing neurons in the Arc of mice at 12 months of age or in the Arc of postmortem hypothalamus of normal human donors. (**C**) Polymerase chain reaction (PCR) analyses of genomic DNA, (**D**) qPCR analyses of RNA, and (**E**) immunoblotting of membrane protein extracted from the hypothalamus, pituitary gland (Pit), and/or other tissues of 2-3 weeks old male ^Neuron^CaSR^+/−^ (Het), ^Neuron^CaSR^-/-^ mice (KO), and control (Cont) mice. Deletion of CaSR gene (ΔExon7), indicated by a 284-bp DNA fragment in panel C, occurred mainly in all brain regions, but not in the Pit or other peripheral tissues. Weak signals of CaSR gene in lung, intestine, and kidney were likely due to inclusion of minute neuronal elements in the tissues. N=3 mice per group. Tagman-based qPCR analyses with a primer set flanking the junction of exon 6 and 7 in panel D, confirmed the ablation (>95%) of CaSR RNA in the hypothalamus but not in the Pit of the KO mice. RNA expression levels were normalized to the expression of ribosomal protein gene L19. Mean ± SEM ****p<0.0001 by 1-way ANOVA N= 6-8 mice per group. A representative immunoblotting of membrane protein using antisera against the C-terminal tail of the CaSR in panel E showed the ablation of the glycosylated receptor proteins (140 and 160 kD bands) in the Hyp, but not in the Pit, of the KO mice. N=3-6 mice per group.

Quantitative real-time PCR (qPCR) analyses showed a >95% reduction in CaSR mRNA level in the hypothalamus (**Fig. 1D, Hyp-CaSR6/7)** of ^Neuron^CaSR^-/-^ (**KO**) mice vs control (**Cont**) littermates. We observed a gene dosage effect in suppressing 40% CaSR RNA levels in the hypothalami of heterozygous ^Neuron^CaSR^+/−^ mice (**Fig. 1D, Hyp-CaSR6/7; Het**). There was a ≈30% reduction in CaSR RNA level in the pituitary of the KO mice, likely due to the loss of CaSR RNA in the posterior pituitary, largely composed of neuronal projections from the hypothalamus (**Fig. 1D, Pit-CaSR6/7**). Immunoblotting confirmed the ablation of CaSR protein in the hypothalamus of ^Neuron^CaSR^-/-^ (**KO**) mice (**Fig. 1E, Hyp)**. In contrast, CaSR protein levels were somewhat increased in the pituitary of KO mice (**Fig. 1E, Pit**), indicating potential compensatory feedback.

### Neuronal knockout of CaSR results in major growth and skeletal defects

Reduced body sizes and weights (by up to ≈25%) were apparent in ^Neuron^CaSR^-/-^ mice at 2 weeks and 3 months of age (MOA) (**Fig. 2A,B**) and persisted throughout life (data not shown). 3D- reconstructed micro-computed tomography (µCT) images demonstrated decreased femur length (**Fig. 2C, left panel**) and trabecular bone mass (**Fig. 2C, right panel**) in the KO vs Cont littermates. Structural analyses of distal femurs showed significantly reduced trabecular tissue volume (Tb.TV), bone fraction (Tb.BV/TV), and number (Tb.N), unchanged thickness (Tb.Th) and connectivity density (Tb.CD), and/or increased trabecular spacing (Tb.Sp) in the KO vs Cont bone (**Extended Data Fig. 2A,B)**. Similarly, we observed decreased cortical tissue volume (Ct.TV) and bone volume (Ct.BV) at the tibiofibular junctions of the KO vs Cont mice, indicating a general delay in skeletal development in the KO mice (**Extended Data Fig. 2A,B)**.

**Fig. 2.**
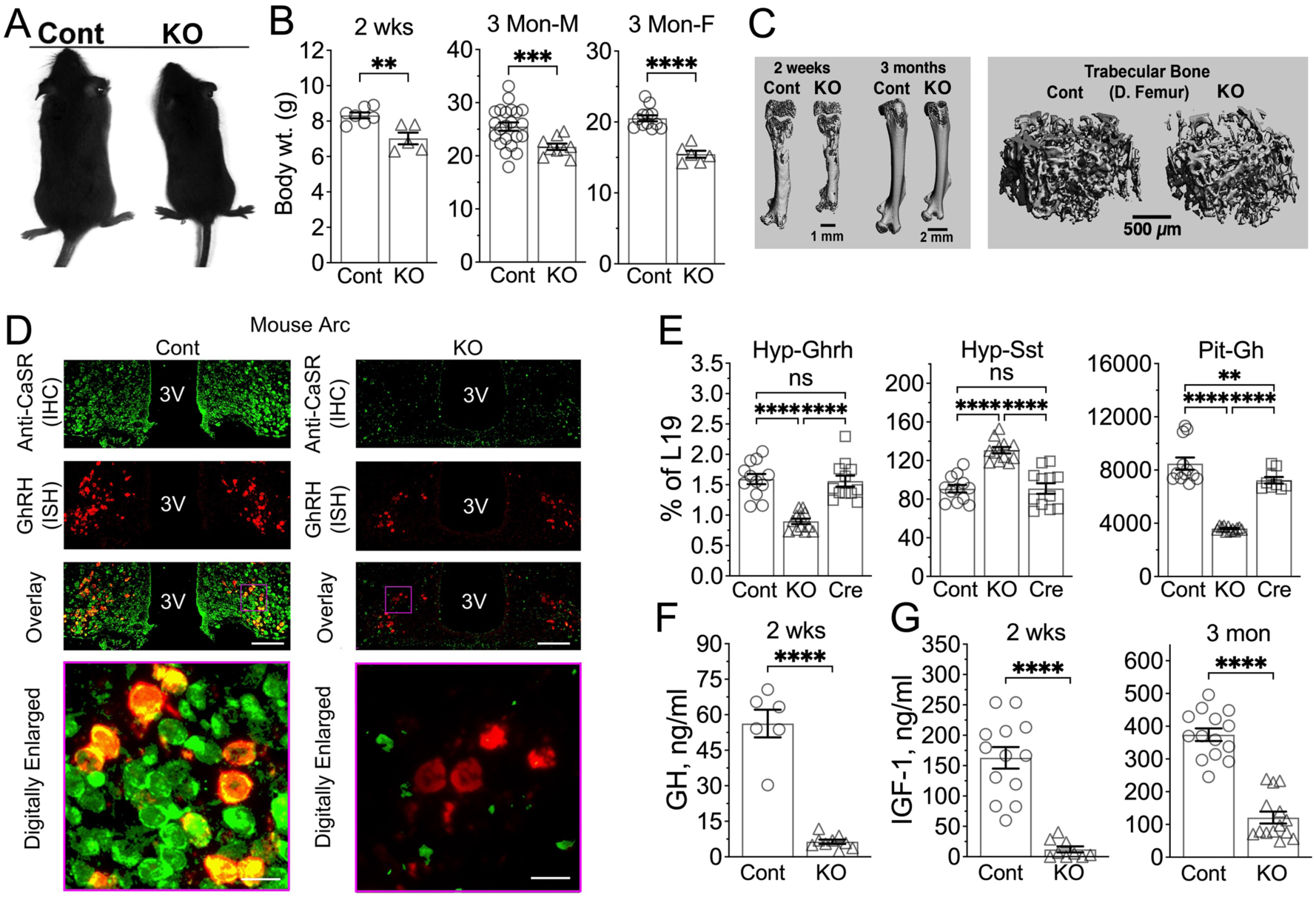
Delayed growth and skeletal development due to dysregulated Hyp-Pit-GH/IGF1 axis in the ^Neuron^CaSR^-/-^ mice. (**A,B**) A negative impact of neuronal *Casr* gene KO on body growth was revealed by (**A**) photographs of 3-month-old male mice and (**B**) body weights (wt.) of 2-weeks-old male and 3-months-old male and female mice. **p<0.01, ***p<0.005, and ****p<0.0001 by 2-tailed Student t-test N=5-14 mice per group. (**C**) Delayed skeletal development in male KO versus Cont mice was shown by 3D-reconstructed µCT images of 2-week and 3-month-old whole femurs (left panel) and trabecular bone elements in the secondary spongiosa of the distal femur (right panel). (**D**) Concurrent IHC and ISH detections show ablation of CaSR protein (in green) along with reduced expression of GHRH RNA (in red) in Arc of 12-month-old ^Neuron^CaSR^-/-^ mice versus Cont littermates. N=4 mice/genotype. Scale Bar = 125 µm and 20 µm. (**E**) qPCR analyses of extracted RNA confirmed quantitatively the reduced GhRH RNA and increased somatostatin (SST) RNA expression in the hypothalamus (Hyp) and reduced growth hormone (GH) RNA expression in the Pit of ^Neuron^CaSR^-/-^ mice versus 2 strains of age-matched control littermates -- CaSR^fl/fl^ (Cont) and Nestin-Cre (Cre) mice. RNA levels were normalized to the expression of L19. **p<0.01 N= 6-8 mice per group. (**F**) Serum GH and (**G**) IGF1 levels in 2 weeks and/or 3 months old ^Neuron^CaSR^-/-^ mice were reduced by >90% and >80%, respectively, compared to the levels in CaSR^fl/fl^ control littermates (Cont). **p<0.01 and ****p<0.0001 by 2-tailed Student t-test N=6-14 mice.

In ^Neuron^CaSR^-/-^ humeri, the expression of mature osteoblast marker osteocalcin (Ocn or Bglap), and mineralizing enzymes ankyrin 1 (Ank1) and ectonucleotide pyrophosphatase/phosphodiesterase 1 (Enpp1) were significantly decreased (**Extended Data Fig. 2C)**, whereas early immature osteoblast marker type I collagen [α_1_(I) encoded by Col1a1] was increased (**Extended Data Fig. 2C)**. These data are consistent with a delay in osteoblast differentiation and a dysregulated mineralizing function in the bone of KO mice. These skeletal defects were not due to changes in the level of CaSR expression in osteoblasts, as this was not altered **(Extended Data Fig. 2C**). There was, however, a significant reduction in IGF1 expression in the KO vs control bones, suggesting that impaired IGF1 signaling in bone may contribute to the skeletal phenotype. Interestingly, the expression of IGF1R RNA was modestly elevated in the KO bone (**Extended Data Fig. 2C**), perhaps representing a compensatory effect. In the ^Neuron^CaSR^-/-^ mice, serum Ca^2+^ (Cont: 10.16 ± 0.19; KO: 10.32 ± 0.17 mg/dL) and Pi (Cont: 6.14 ± 0.20; KO: 6.06 ± 0.32 mg/dL) levels were unchanged (data not shown), excluding the development of hypercalcemia or hyperparathyroidism as the cause of their skeletal defects.

### Mice lacking neuronal CaSR have an impaired hypothalamo-pituitary GH/IGF1 axis

The impaired IGF1 signaling in the bone of ^Neuron^CaSR^-/-^ mice led us to hypothesize that neuronal CaSR KO may interfere with the GH/IGF1 neuroendocrine axis. Indeed, hypothalamic expression of GHRH RNA was reduced by 45% (**Figs. 2D, 2E Hyp:GHRH**), while the hypothalamic expression of somatostatin, an inhibitor of GH secretion (**Fig. 2E Hyp:SST**), was increased by 30% in 2-week-old KO mice vs. either CaSR^fl/fl^ (Cont) or Nestin-Cre (Cre) control littermates. Pituitary GH mRNA levels were reduced by 60% in KO mice **(Fig. 2E, Pit:GH).** Consistent with this, serum growth hormone (GH) **(Fig. 2F)** and IGF1 **(Fig. 2G)** levels in the KO mice were markedly reduced. These data support an essential role for neuronal CaSR in the hypothalamo-pituitary GhRH/GH/IGF1 axis.

The expression of TRH, CRH, and GnRH was also significantly decreased in ^Neuron^CaSR^-/-^ hypothalami **(Extended Data Fig. 3A**), and in accord with this, expression of their downstream anterior pituitary hormones was also lower (**Extended Data Fig. 3B**). In contrast, hypothalamic oxytocin (OXT) and pituitary prolactin (PRL) expression, which are under tonic inhibition, were increased in the ^Neuron^CaSR^-/-^ mice. These data indicate that CaSR ablation suppressed hypothalamic function leading to panhypopituitarism.

### Neuronal knockout of CaSR drives dysregulated energy metabolism

At 3 months of age, ^Neuron^CaSR^-/-^ mice showed an increase in visceral fat **(Fig. 3A)**, with a 60% increase in the combined weights of retroperitoneal, peri-gonadal, and peri-nephric fat pads and a 25% increase in total body fat content by DEXA scan (**Fig. 3B,C)**. The increased adiposity was reflected in increased serum leptin **(Fig. 3D**). The KO mice also showed significantly poorer glucose tolerance in an intraperitoneal glucose tolerance test (IPGTT) **(Fig. 3E)**.

**Fig. 3.**
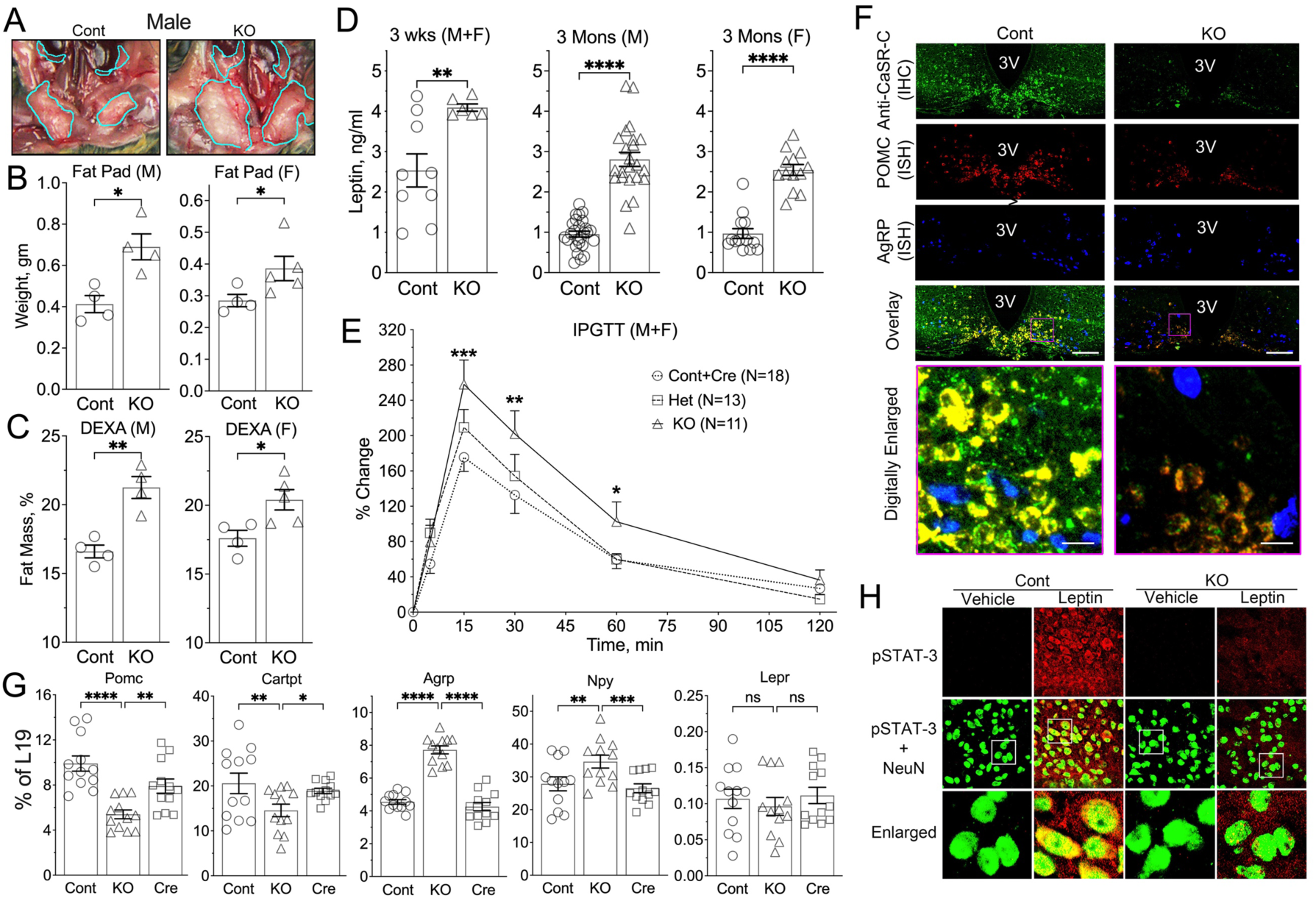
^Neuron^CaSR^-/-^ mice develop obesity. (**A**) Pictures of abdominal cavities with visceral fat contents traced by light-blue lines in 3-month-old male **^Neuron^CaR^-/-^ (**KO) versus Cont mice, (**B**) combined weights of retroperitoneal, peri-gonadal, and peri-nephric fat pads, and (**C**) DEXA measurements of total body fat compositions showed obese phenotypes in 3-month-old male (M) and female (F) KO versus Cont mice. *p<0.05, **p<0.01, and ****p<0.001 by 2-tailed Student t-test N= 4-12 mice per group. (**D**) Serum leptin levels increased in 3 weeks and 3 months old KO versus Cont mice determined by ELISA. **p<0.05, ****p<0.0001 N=6-14 mice. (**E**) IPGTT assessed by measuring tail-vein blood glucose levels following IP injections of glucose (2mg/g body wt.) and presented as % change from the basal levels [121 ± 5.0, 116 ± 4.6, and 101 ± 4.2 mg/dl in Cont, heterozygous (Het), and homozygous (KO) KO mice respectively) revealed glucose intolerance in the mixed male/female KO versus Het and Cont mice. *p <0.05, **p<0.01, and ***p<0.005 by 2-way ANOVA N=11-18 mice per group. **(F)** Concurrent IHC and ISH assays show ablation of CaSR protein (in green) along with reduced POMC RNA expression (in red) and increased ArgRP RNA expression (in blue), in sections of Arc nucleus from ^Neuron^CaSR^-/-^ mice versus Cont littermates. N=4 mice/genotype. Scale Bar = 125 µm and 20 µm. **(G)** qPCR analyses of extracted RNA showed decreased expression of *Pomc* and *Cartpt* genes and increased expression of *Agrp* and *Npy* genes without altering *Lepr* expression in the hypothalamus of 2 weeks old ^Neuron^CaR^-/-^ (KO) versus Cont and Cre mice. *p<0.05, **p<0.01 ***p<0.005, and ****p<0.0001 by 1-way ANOVA, N=8-12 mice per group. (**h**) Florescent IHC showed increased expression and colocalization of phosphorylated STAT3 (in red) with neuronal nuclear protein, NeuN (in green) in the Arc of 3-months-old ^Neuron^CaSR^-/-^ versus and control (Cont) mice following ICV injections of leptin (Lepn, 1.5 ng in 3 µl PBS) versus vehicle (Veh). Brain samples were harvested and processed 10 min after injection. White boxes inside the images in the middle row depict the areas to be enlarged and shown in the bottom row. N = 4 mice per group mice.

To investigate whether the imbalanced energy metabolism in ^Neuron^CaSR^-/-^ might be due to changes in the responses of POMC and AgRP neurons to leptin, we examined the hypothalamic expression of neuropeptides expressed in these neurons. Despite a significant increase in serum leptin levels and unaltered RNA expression of LEPR, ^Neuron^CaSR^-/-^ hypothalami showed a 30% reduction in POMC and CART expression, while AgRP and NPY expression was increased by 70% and 25%, respectively (**Fig. 3F, 3G**). In control mice, a single ICV injection of leptin profoundly stimulated the phosphorylation (an activation surrogate) and nuclear translocation of STAT-3 in hypothalamic neurons (**Fig. 3H-Cont**), but these responses were suppressed in the KO (**Fig. 3H-KO**), indicating impaired leptin responsiveness. Together, these data suggest that mice lacking neuronal CaSR have reduced central leptin sensitivity.

### Knockdown of CaSR specifically in the arcuate nucleus disrupts energy and skeletal homeostasis

To investigate whether the effects of neuronal CaSR knockout were mainly driven by its loss in the Arc, we generated mice in which CaSR was specifically knocked down in the Arc (^ARC^CaSR^-/-^). CaSR floxed mice were bilaterally injected into the Arc with AAV2-eGFP-Cre or control AAV2- eGFP viral particles. ^ARC^CaSR^-/-^mice had a ≈50% reduction in CaSR expression in the Arc, but unchanged CaSR expression in the nearby ventromedial nucleus (VMN) (**Fig. 4A)**, characterized by much lower expression of AgRP, NPY, and PMOC, and other Arc-specific cell markers (**Extended Data Fig. 4A**). ^ARC^CaSR^-/-^mice had increased body weight and adiposity by EchoMRI (**Fig. 4B,C**), reduced glucose tolerance (**Fig. 4D**) and decreased sensitivity to the effects of peripherally administered leptin on food intake compared to controls (**Fig. 4E**). As anticipated, Arc of ^ARC^CaSR^-/-^ showed reduced CaSR, POMC and Ghrh RNA levels (**Fig 4F and Extended Data Fig. 4B)** and altered expression of genes associated with reduced insulin/leptin signaling (**Fig. 4G**), despite an increased leptin level (**Fig. 4H**). A strong correlation between POMC and CaSR RNA expression (**Fig. 4I**), supports a close regulation of POMC expression by CaSR signaling.

**Fig. 4.**
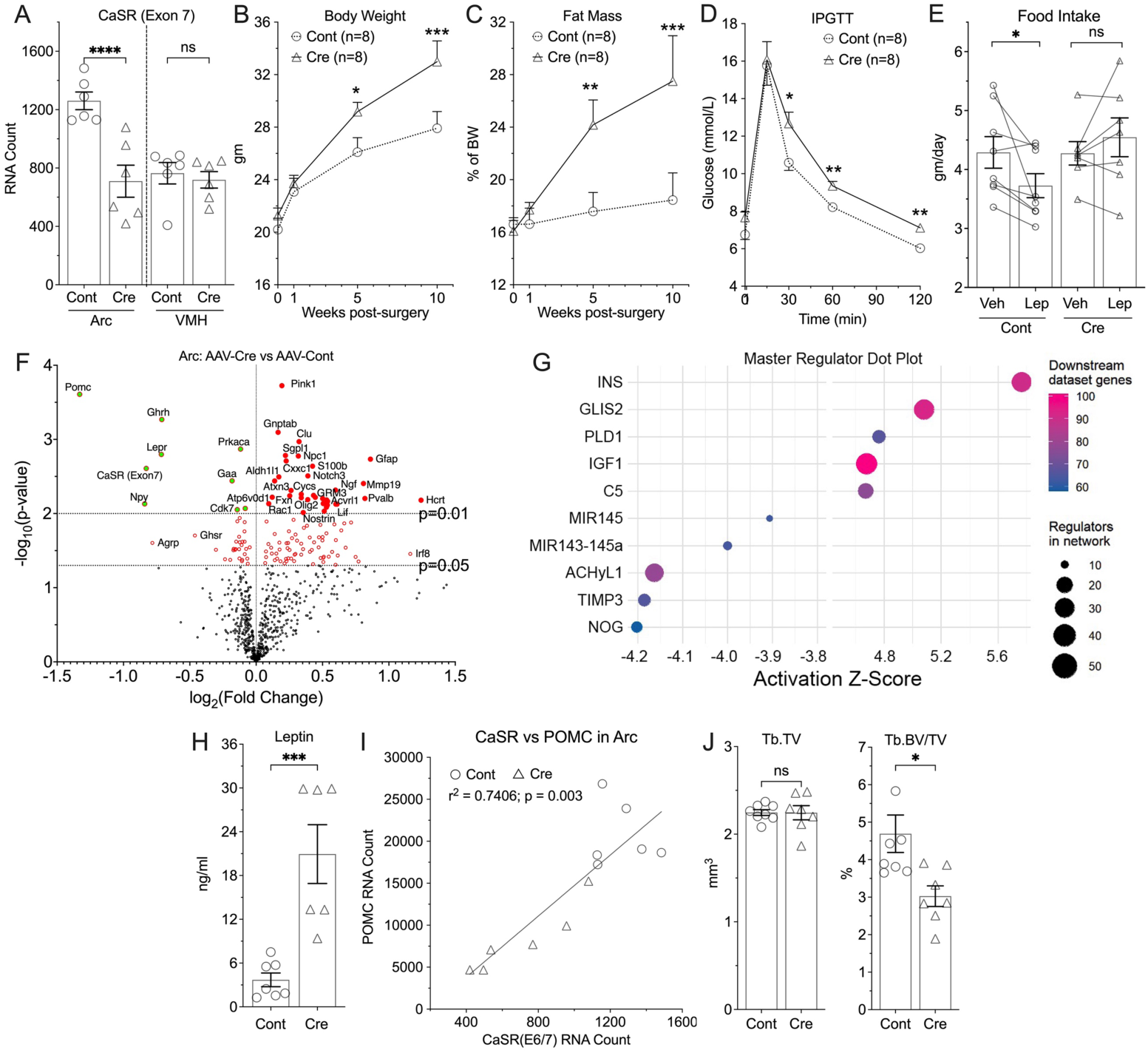
^ARC^CaSR^-/-^ mice have increased fat mass and reduced leptin sensitivity. (**A**) nCounter analyses of extracted RNA from micro-dissected Arc and VMH of 8-12 weeks mixed male and female CaSR^flox/flox^ mice injected with AAV2-eGFP-Cre (^ARC^CaSR^-/-^) or AAV2-eGFP (Cont) to knock down CaSR expression in the Arc. Mean ± SEM. ****p<0.0001 by 2-tailed t-test, N=6-12 mice. Measurements of (**B**) body weight, (**C**) fat mass, (**D**) IPGTT response, and (**E**) 24-hours food-intake in response to single injection of leptin in the ^ARC^CaSR^-/-^ versus Cont mice. (**F**) A volcano plot and (**G**) pathway analyses of RNA profiles, (**H**) plasma leptin levels, and (I) a linear correlation between CaSR and POMC RNA expression in the Arc of ^ARC^CaSR^-/-^ and Cont mice support a close interaction between these 2 signaling cascades in the anorectic neurons. (J) Tb size (Tb.TV) and fractions (Tb.BV/TV) in the ^ARC^CaSR^-/-^ versus Cont mice. Mean ± SEM *p<0.05, **p<0.01, ***p<0.005, and ****p<0.0001 by 2-tailed t-test, N=6-7 mice per group.

Ingenuity Pathway Analysis IPA (http://www.ingenuity.com/) was conducted on the nCounter expression data for 800 genes using a cutoff p value of <0.05 which reduced the analysis ready dataset to 210 genes, among which 152 genes were upregulated and 58 were downregulated. The analysis predicted that likely upstream master regulators of these changes included insulin (INS) (ranked highest with a z-score of 5.8, 91 downstream dataset genes) and IGF-1 (z-score 4.7, 101 downstream dataset genes)(**Fig. 4G**). Using IPA upstream regulator analysis to identify transcriptional regulators that might account for the observed gene changes in our dataset, insulin (INS) was predicted to drive a mechanistic network to modulate a number of genes associated with metabolic regulation in the arcuate nucleus (**Extended Data Fig. 4C**). ^ARC^CaSR^-/-^ mice also had a bone phenotype, albeit less pronounced than that of ^Neuron^CaSR^-/-^ mice, with reduced Tb fraction without altered bone sizes (**Fig 4J**).

The role of the CaSR in systemic Ca^2+^ regulation is well characterized, but relatively little attention has been given to its role in growth and energy metabolism. However, it is broadly expressed in tissues vital to those functions, including the hypothalamus^27,28^, adipocytes^29^, and pancreatic beta- cells^30^. CaSR expression in specific hypothalamic neuronal populations and the smaller body size, glucose intolerance, increased adiposity, delayed bone development and mineralization, and panhypopituitarism observed in the ^Neuron^CaSR^-/-^ mice provide compelling *in vivo* evidence for previously unidentified actions of neuronal CaSR in supporting growth, energy, and skeletal metabolism. These neuroendocrine actions are independent of the actions of CaSR in PTCs and osteoblasts, as its expression in PTCs remained unchanged and serum PTH and Ca^2+^ levels were unaltered in the ^Neuron^CaSR^-/-^ mice. Also, in contrast to the early death (within 3-4 weeks of birth) of the osteoblast- and PTC-specific KO mice^3^, ^Neuron^CaSR^-/-^ mice were viable postnatally despite their apparent growth and metabolic derangements.

Our observations in the ^Neuron^CaSR^-/-^ mice are supported by previous *in vitro* and *in vivo* observations demonstrating various CaSR functions in neurons. CaSR might function as a regulator of neuronal migration during development^31^ and newborn ^Global^CaSR^-/-^ mice showed lower brain weight and size, and decreased expression of neuronal and glial differentiation markers, suggesting CaSR plays a critical role in neuronal development^32^. However, the central effects of CaSR are not simply developmental. In cultured neurons, activation of CaSR stimulates various ion channels, activates phospholipase C, and increases [Ca^2+^]_i_ in soma^33–35^. At nerve terminals the CaSR regulates membrane excitability^36^ and modulates neurotransmitter release^37^. The body weight and skeletal phenotype of the ^ARC^CaSR^-/-^ mice, in which CaSR was knocked down specifically in the Arc in adult animals, also demonstrates that CaSR in the brain plays roles beyond development, and show the importance of CaSR in the Arc, particularly in energy homeostasis. Interestingly, we did not observe significant growth, skeletal, and metabolic changes when we previously targeted CaSR ablation specifically to hippocampal and cortical neurons^38^. Those data suggest that the role of neuronal CaSR in body growth and skeletal and metabolic homeostasis is specific to brain region and likely to specific neuronal types, such as POMC and GHRH neurons due to their high levels of CaSR expression and strong responses to CaSR KO or knockdown. Further work is required to determine whether the differences between the ^ARC^CaSR^-/-^ and ^Neuron^CaSR^-/-^ models reflect the higher level of knockdown in the arcuate nucleus in the ^Neuron^CaSR^-/-^ mouse, or whether some of the greater effects on bone and the growth axis are mediated by pathways upstream of the arcuate nucleus.

Leptin alters feeding behavior and neuroendocrine function in the hypothalamus, by binding to LEPR and activating intracellular Jak2/STAT-3 signaling responses^39^. ICV leptin injections increase GHRH and POMC RNA levels and suppress AgRP expression in rodents^40,41^. Our observation of the profoundly reduced ability of ICV leptin injection to increase phosphorylation and nuclear translocation of STAT-3 in the hypothalamic neurons of ^Neuron^CaSR^-/-^ mice suggest CaSR signaling is critical in supporting a full neuronal response to leptin. This is also consistent with the decreased GHRH and POMC mRNA expression and increased AgRP and NPY levels in the hypothalami of ^Neuron^CaSR^-/-^ mice in the presence of a high serum leptin level. We ascribed this leptin insensitivity to the general hypothalamic dysregulation that leads to growth retardation, glucose intolerance, and increased intrabdominal fat mass seen in the ^Neuron^CaSR^-/-^ mice. Similarly, the apparent skeletal defects in the ^Neuron^CaSR^-/-^ mice could be attributed to the dysregulated hypothalamo-pituitary GH/IGF1 axis, as demonstrated by reduced IGF1 RNA expression in the bone of ^Neuron^CaSR^-/-^ mice. Alternatively, the increased leptin levels in the mice may also be a contributing factor^24^, as leptin has been demonstrated to be a direct regulator of bone mass by inhibiting osteoblastic bone formation ^42^.

In summary, we present evidence for critical actions of the CaSR in controlling growth, skeletal, and energy metabolism by modulating neuroendocrine functions in hypothalamic neurons. Given the importance of the role of the hypothalamus in growth, energy homeostasis and appetite, future studies should focus on the role of CaSR in specific subpopulations of hypothalamic neurons (e.g., GHRH, POMC, and AgRP). In particular, work elucidating the interplay between LEPR and CaSR signal responses is needed to understand the precise mechanisms that critically integrate mineral, skeletal, and energy metabolism to ensure adequate proportions of soft and hard tissue development and support normal physiological functions in the body.

## Materials and Methods

### Generation and genotyping of mouse models

All mice were kept in a climate-controlled room with a 12-hour light/12-hour dark cycle as previously described ^3^ with ad libitum access to food and water, unless otherwise stated. All protocols performed in the US were approved by the Institutional Animal Care and Use Committee of the San Francisco Department of Veterans Affairs Medical Center. Experiments conducted in the UK were performed under authority of the U.K Animals (Scientific Procedures) Act, 1986, and approved by the Animal Welfare Ethical Review Body of Imperial College London.

*In vivo* ablation of the CaSR in neurons was achieved by mating floxed-CaSR (CaSR^fl/fl^) mice, which carry loxP sequences flanking the exon 7 of the *Casr* gene^3^, with Nestin-Cre mice ^26^. We compared phenotypes of heterozygous (^Neuron^CaSR^+/−^ or Het) and homozygous (^Neuron^CaSR^-/-^ or KO) CaSR knockout mice with 2 control mouse strains: (1) CaSR^fl/fl^ (Cont) mice, which carry homozygous floxed-CaSR alleles without Cre expression, and (2) Nestin-Cre (Cre) mice, which express Cre but not floxed-CaSR alleles. Because the effects of CaSR KO were comparable in both male and female mice, only data from male mice are reported.

Genotyping of mice was performed by PCR analysis of genomic DNA extracted from tail snips using primer sets as described ^3,26,43–45^. To detect the presence of the exon7-less *Casr* allele, genomic DNA from different tissues of ^Neuron^CaSR^-/-^ were analyzed by PCR with a specific primer set (upper primer, 5’-CCTCGAACATGAACAACTTAATTCGG-3’; lower primer, 5’- CGAGTACAGGCTTTGATGC-3’)^3^.

### Arcuate nucleus CaSR knockdown

8-12 week old CaSR floxed mice were bilaterally injected into the Arc with 0.4 µl of AAV2.CMV.PI.EGFP.WPRE.bGH (AAV2-eGFP) or AAV2.CMV.HI.eGFP-Cre.WPRE.SV40 (AAV2-eGFP-Cre) at a rate of 0.2 µl/min (stereotaxic coordinates relative to bregma: anteroposterior −1.2mm; mediolateral +/− 0.2mm, dorsoventral −5.3mm). Plasmids were a gift from James M Wilson and viral vectors were purchased from Addgene (pAAV.CMV.PI.EGFP.WPRE.bGH; Addgene viral prep # 105530-AAV2) (pAAV.CMV.HI.eGFP-Cre.WPRE.SV40; Addgene viral prep # 105545-AAV2).

### Intracerebroventricular (ICV) leptin injection

ICV leptin injections were carried out in 3 month old ^Neuron^CaSR^-/-^ and Cont littermates as described previously^46^. Briefly, 3 µl of recombinant mouse leptin stock (0.5 ng/µl PBS; Cat#498-OB, R&D Systems) or vehicle (PBS) was injected into the right lateral cerebral ventricle (stereotaxic coordinates: 1 mm caudal to bregma, 1.3 mm lateral to sagittal suture, and 2 mm in depth) at a speed of 0.3 µl/min via a burr hole. Brains were removed 10 min after the injections for analyses.

### Leptin Food Intake Studies

Individually housed, *ad libitum* fed, ^ARC^CaSR^-/-^ mice were intraperitoneally injected with 1mg/kg mouse recombinant leptin (kindly provided by A. F. Parlow at the ‘National Hormone & Peptide Program’) at the onset of the dark phase. Food intake was measured at 24 hrs. post administration.

### Immunohistochemistry (IHC), *in situ* hybridization (ISH), and Immunoblotting

To localize CaSR expression in specific populations of hypothalamic neurons, formalin-fixed paraffin-embedded (FFPE) brain sections from 12-month-old ^Neuron^CaSR^-/-^ and Cont littermates and 5 human subjects with low AD neuropathological changes (2 females and 3 males at Braak stages 1 or 2, 63-76 years of age at death, as prepared in the laboratory of Dr. Lea Grinbergs, co-Leader of the UCSF/Neurodegenerative Disease Brain Bank, at UCSF Weill Institute for Neurosciences) were subjected to concurrent IHC detection of CASR protein with Texas red-conjugated rabbit anti-CaSR-N antibody (10 μg/ml)^47^ and ISH detection of mouse or human POMC, AgRP, and/or GhRH RNA using AF488 or Cy5-conjugated RNAScope probes (ACD, Newark, CA) according to manufacturer’s instructions. Tissue images were acquired, pseudo-colored, and overlayed. Specificity of the antibody was confirmed by the absence of signal in sections from ^Neuron^CaSR^-/-^ mice or sections subjected to non-immune IgGs.

Brain sections from mice with ICV-injected leptin or vehicle were co-labeled with anti-rabbit pSTAT-3 (1:200, Cell Signaling) plus anti-mouse NeuN (1:200, Chemicon) and corresponding Alexa Fluor 488-conjugated goat anti-mouse IgG (1:200, Invitrogen) and Alexa Fluor 594- conjugated goat anti-rabbit IgG (1:200, Invitrogen).

To semi-quantify CaSR protein expression, crude plasma membranes were prepared from hypothalami and pituitaries of 2 week-old mice, followed by protein extraction with a RIPA buffer as previously described^48^. Protein samples (50 µg) were electrophoresed and immunoblotted for CaSR and ß-actin (as loading control) using an anti-CaSR antibody (1 µg/ml) and anti-ß-actin (0.2 µg/,l) and an HPP-conjugated anti-rabbit IgG antibody (2 µg/ml) as described^3^.

### Serum chemistries

Blood was drawn by cardiac puncture after euthanasia by isoflurane inhalation, followed by tissue harvest. Serum samples were prepared by centrifugation and analyzed for total serum calcium using a bioanalyzer, and intact PTH, GH, IGF1, and leptin using commercial PTH (Immutopics, San Clemente, Ca, USA), GH, IGF1, and leptin ELISA kits (R&D Systems, Minneapolis, MN, USA), respectively, following manufacturer instructions.

### Analyses for gene expression

#### Quantitative PCR (qPCR)

Total RNA samples isolated from the hypothalamus, pituitary and bone (without marrow) of 2 week-old ^Neuron^CaSR^+/−^, ^Neuron^CaSR^-/-^, and control mice were reverse transcribed by M-MLV reverse transcriptase into cDNA and subjected to TaqMan-based qPCR^3^ and for gene expression profiles using commercial sets of primers and probes (Applied Biosystems). Expression levels of genes of interest were presented as the percentage of level of the mitochondrial ribosomal protein L19.

#### nCounter ensemble RNA analyses PCR

Total RNA samples (250 ng) isolated from micro- dissected Arc and VMN of ^ARC^CaSR^-/-^ and control mice were subjected to the nCounter assay. Briefly, the RNA samples were hybridized with specific probes conjugated with 4-color, 6-spot optical barcodes, washed, and digitally counted using the nCounter system (Bruker Spatial Biology, Inc., Seattle, WA). These probes include a commercially available (Mouse Neuropathology) panel containing 760 genes and a spike-in probe set of 40 genes. Nanostring nSolver software was used to normalize the digital counts using internal spike-in controls (ERCCs) and a housekeeping gene panel as previously described (71). Gene expression data from the ARC of ^ARC^CaSR^-/-^ and control mice were compared using Ingenuity Pathway Analysis version 134816949 (Qiagen). Reads were mapped to mouse GRCm38 genome and genes quantitated using STAR (2.5.0a) using default parameters. Differential gene expression was determined using gene-wise likelihood ratio tests from EdgeR (3.34.0)/R 4.1.0, with significance threshold set at uncorrected p<0.05.

### Skeletal analyses by micro-computed tomography (µCT)

To compare bone mineral content and structural parameters, we performed micro–computed tomography (μCT) scans at two sites: distal femur for trabecular (Tb) bone and tibiofibular junction (TFJ) for cortical (Ct) bone as described^3^. Briefly, femurs and tibias were isolated, fixed in 10% Neutral Buffered Formalin (PBF) for 24 hours, stored in 70% ethanol, and scanned by a SCANCO viva CT 40 scanner (SCANCO Medical AG, Basserdorf, Switzerland) with 10.5 µm voxel size and 55-kV X-ray energy. A threshold of 420 mg hydroxyapatite (HA)/mm^3^ was applied to segment total mineralized bone matrix from soft tissue. 3-D image reconstructions and analyses were performed using the manufacturer’s software to obtain the following structural parameters.

### Assessment of adiposity

Whole-body DEXA scans were performed under general anesthesia with isofluorane (1%) inhalation using a Lunar Piximus2 (GE Lunar, Waukesha, WI) to determine whole body fat and lean body compositions. Retroperitoneal, peri-gonadal, and peri-nephric fat pads were dissected from ^Neuron^CaSR^-/-^ and control littermates and weighed. Combined fat weights were presented and compared.

Lean and fat mass, expressed as a percentage of total body weight, were assessed in restrained ^ARC^CaSR^-/-^ and control mice by nuclear magnetic resonance imaging using an Echo MRI-100H body composition analyzer (Echo medical systems, Huston, Texas) Measurements were acquired 0, 5 and 10 weeks post-surgery, using triple acquisition settings.

### Intraperitoneal Glucose Tolerance Testing (IPGTT)

^Neuron^CaSR^-/-^ and ^ARC^CaSR^-/-^ and controls were fasted overnight or for 6 hours with free access to water, respectively. Tail vein blood was sampled immediately before glucose injection to determine the basal glucose level using a glucose meter. Following the injection of glucose (2mg/g body weight) through an intraperitoneal route, glucose levels from tail vein blood were measured at 15, 30, 60 and 120 minutes post-injection.

### Statistics

Data from two groups were represented as mean ± standard error of the mean (SEM) and compared using unpaired 2-tailed Student’s *t*-test or 1- or 2-way ANOVA. Significance was assigned for *p-value*<0.05.

## Acknowledgements

Grant Support: This work was supported by grants from: Department of Veterans Affairs (1IK6BX004835 and 1I01 BX005851 to WC); NIH (RF1AG075742 and P30AR075055 to WC) and NIDDK (5T32DK007418-35 to JP); and the Endocrine Fellows Foundation (Fellows Development Research Grant Program to JP).

The Section of Endocrinology and Investigative Medicine is funded by grants from the MRC, NIHR and is supported by the NIHR Biomedical Research Centre Funding Scheme and the NIHR/Imperial Clinical Research Facility. KGM is supported by Diabetes UK (18/0005886, 20/0006295), the BBSRC (BB/W001497/1, BB/X017273/1), the MRC (MR/Y013980/1) and the Wellcome Trust (310835/Z/24/Z). The views expressed are those of the author(s) and not necessarily those of DUK, The BBSRC, the MRC, the Wellcome Trust, the NHS, the NIHR or the Department of Health.

## Authors’ roles

Study design: JP, MN, CLT, KGM, and WC. Study conduct: JP, MN, CLT, ZC, NF, HL, SK, AL,, KGM and WC. Data collection: JP, MN, CLT, ZC, NF, HL, SW, AL, KGM, and WC. Data analysis: JP, MN, CLT, GAB, KGM, and WC. Data interpretation: JP, MN, CLT, GAB, KGM, and WC. Draft manuscript: JP, MN, CLT, KGM, and WC. Approving final version of manuscript: JP, MN, CLT, ZC, NF, HL, SK, AL, GAB, KGM, and WC. JP, KGM, and WC take responsibility for the integrity of the data analysis.

---

**Extended Data Fig. 1.**
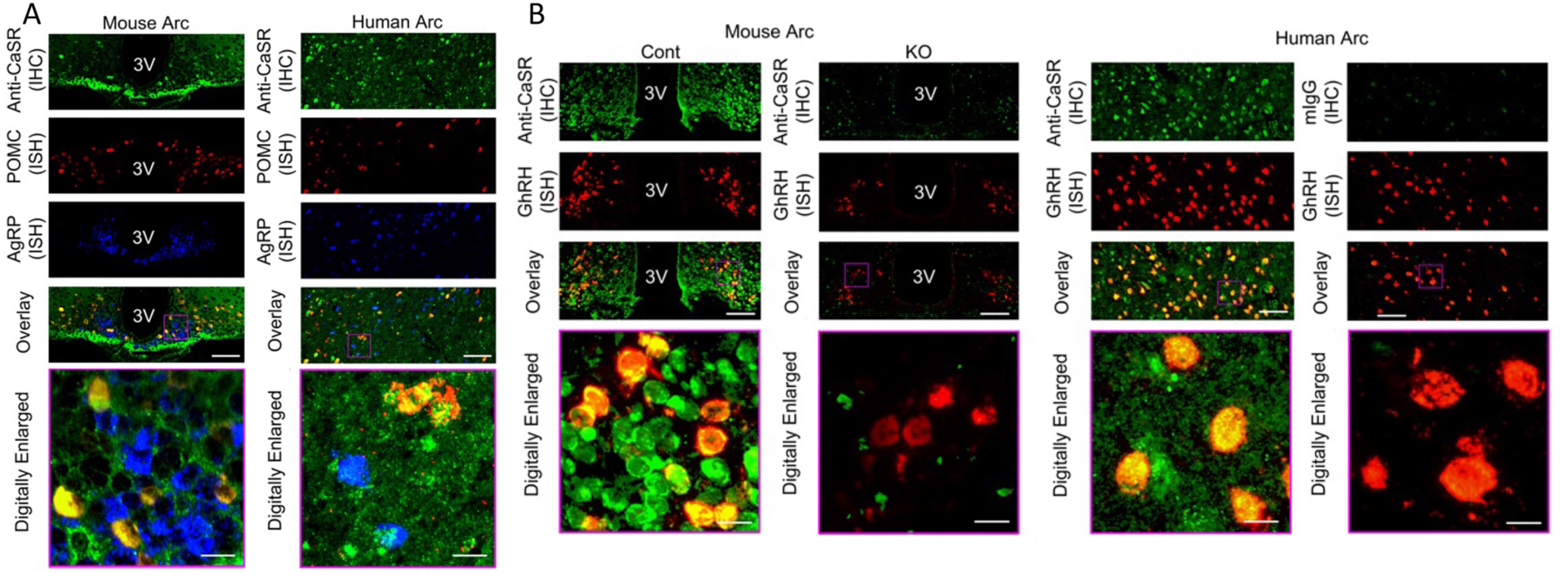
Expression and localization of CaSR protein in neurons expressing POMC or GHRH RNA, but to a lesser extent in the neurons expressing AgRP RNA. (A,B) IHC detection of CaSR (in green) with concurrent ISH detection of POMC (in red) and AgRP (in blue) RNA in the panel A or GHRH RNA in the panel B show strong localization of CaSR in POMC and GHRH neurons (in yellow), but to a lesser extent in AgRP neurons. IHC signals of CaSR were specific due to their absence in the Arc of the ^Neuron^CaSR^-/-^ mice or in human Arc subjected to non-immune mouse IgG. N=3-6 mice per group or 5 human Arc.

**Extended Data Fig. 2.**
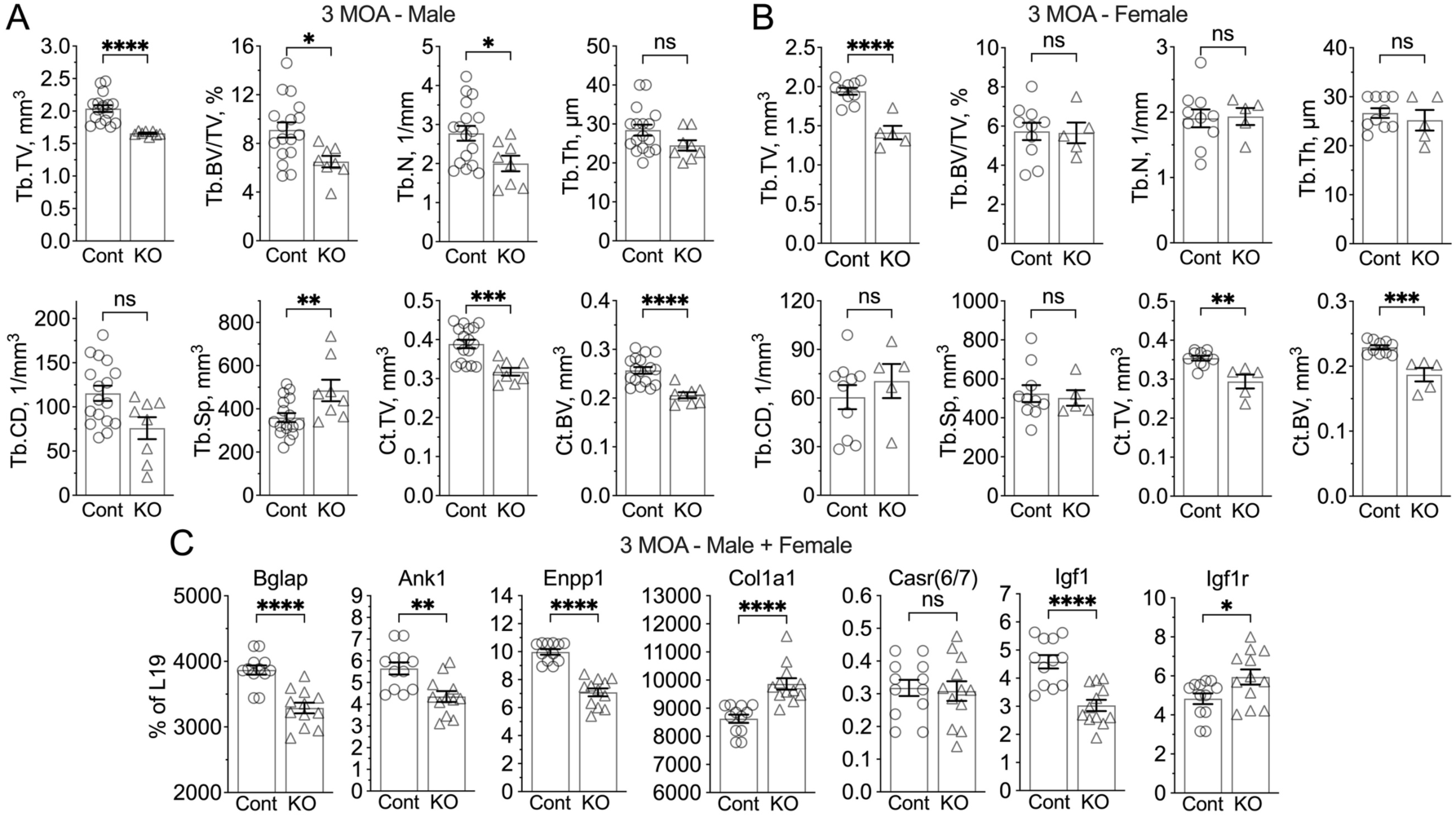
Delayed skeletal development in ^Neuron^CaSR^-/-^ mice. (**A,B**) Quantitation of structural parameters derived from trabecular (Tb) bone in distal femur and cortical (Ct) bone at TFJ of 3-month-old male (A) and female (B) indicate delayed bone development. Mean ± SEM *p<0.05, **p<0.01, ***p<0.005, and ****p<0.0001 by 2-tailed Student t-test N=5-14 mice per group. (**c**) Analyses of RNA extracted from diaphysis of humeri (without bone marrow) of mixed male and female ^Neuron^CaSR^-/-^ mice which show altered expression of differentiation markers [osteocalcin (OCN) and □_1_(I)], mineralization enzymes [ankyrin 1 (ANK1) and ectonucleotide pyrophosphatase/phosphodiesterase family member 1 (ENPP1)], CaSR, IGF1, and IGF1R in osteoblasts/osteocytes. RNA expression levels were normalized to the expression of L19. N= 6 mice per group. *p<0.05, **p<0.01, ***p<0.005, and ****p<0.0001 by 2-tailed Student t-test. ns = not significant. N=5-10 mice per group.

**Extended Data Fig. 3.**
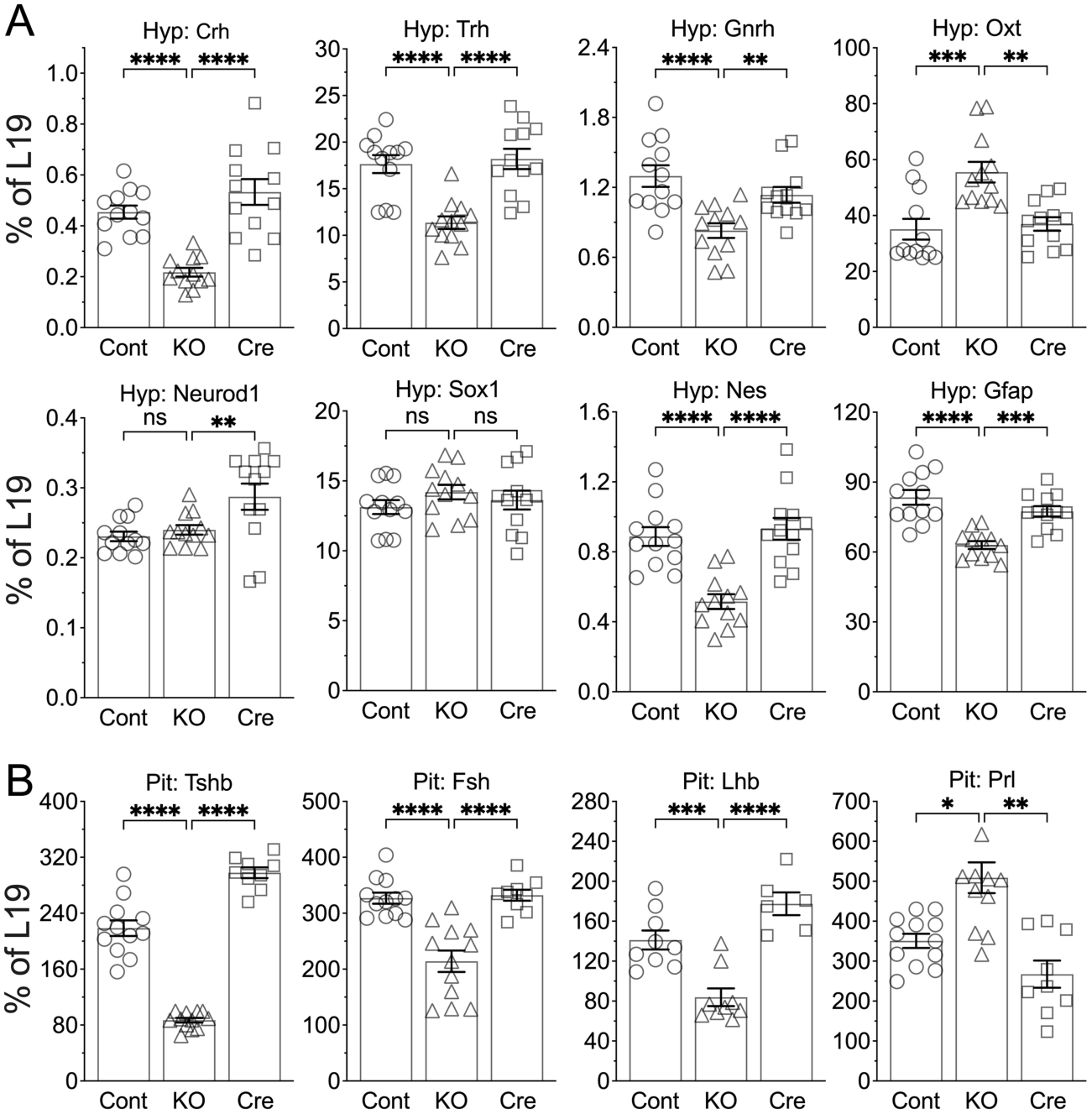
Gene expression profiles showed globally dysregulated hypothalamic functions and panhypopituitarism in the ^Neuron^CaR^-/-^ mice. qPCR analyses were performed to assess expression of each specified gene in the (**A**) Hyp and (**B**) Pit of 2 weeks old mixed male and female ^Neuron^CaR^-/-^ (KO) mice versus Cont and Cre control mice. *p<0.05, **p<0.01, ***p<0.005, and ****p<0.0001. by 1-way ANOVA N=6-12 mice per group.

**Extended Data Fig. 4.**
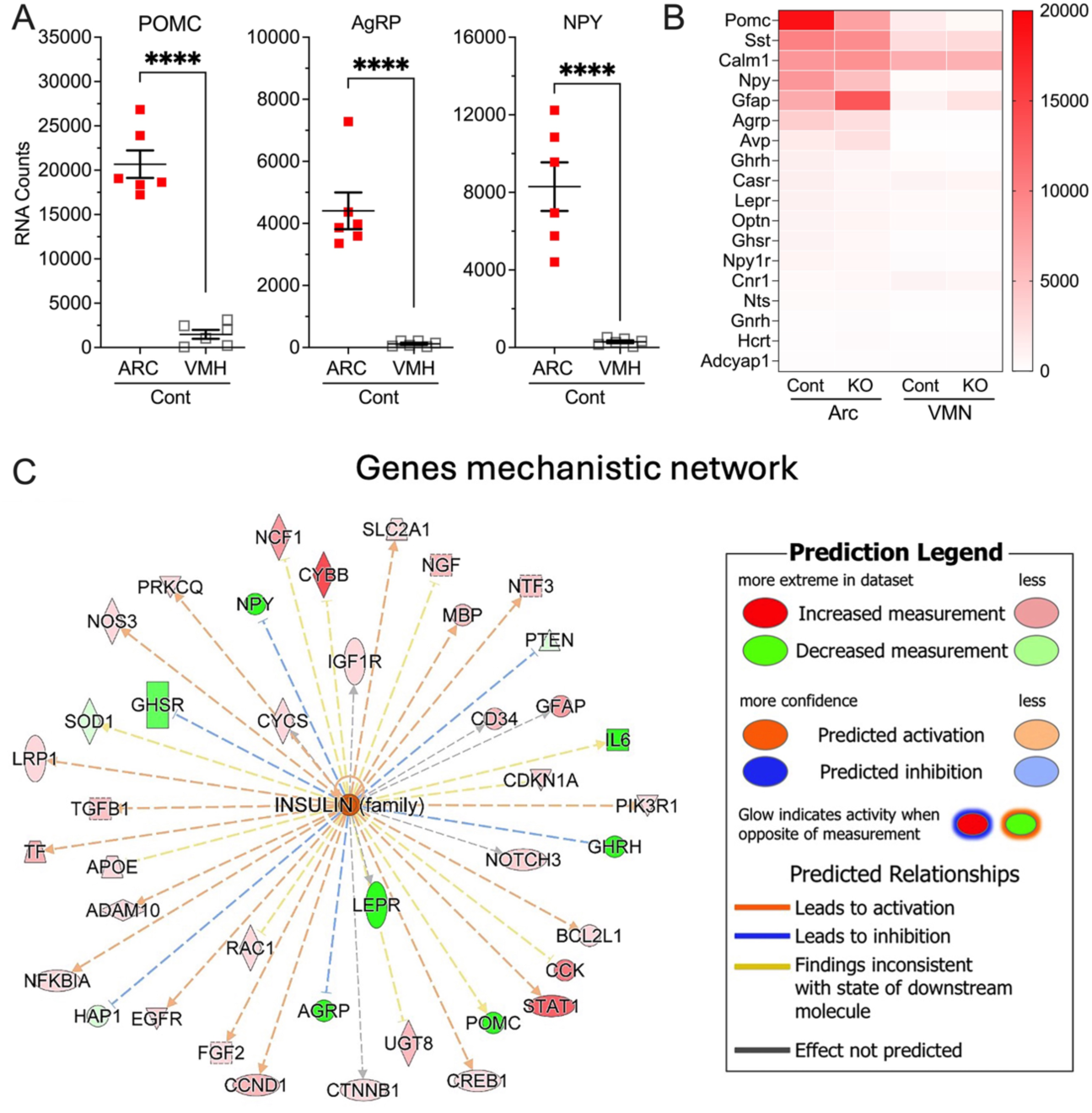
Arc-targeted CaSR knockdown in ^ARC^CaSR^-/-^ mice. (**A**) Dot plot, (**B**) heatmap analyses of RNA counts in micro-dissected Arc and VMH of mice injected with AAV2-eGFP-Cre (^ARC^CaSR^-/-^, KO) or AAV2-eGFP (Cont). Mean ± SEM ****p<0.0001 by 2-tailed t-test, N=6 mice. (**C**) Gene expression data from the ARC of ^ARC^CaSR^-/-^ and control mice were compared using Ingenuity Pathway Analysis version 134816949 (Qiagen). Reads were mapped to mouse GRCm38 genome and genes quantitated using STAR (2.5.0a) using default parameters. Differential gene expression was determined using gene-wise likelihood ratio tests from EdgeR (3.34.0)/R 4.1.0, with significance threshold set at uncorrected p<0.05.

